# Chromosome arm specific patterns of polymorphism associated with chromosomal inversions in the major African malaria vector, Anopheles funestus

**DOI:** 10.1101/068205

**Authors:** Colince Kamdem, Caroline Fouet, Bradley J. White

## Abstract

Chromosomal inversions facilitate local adaptation of beneficial mutations and modulate genetic polymorphism, but the extent of their effects within the genome is still insufficiently understood. The genome of *Anopheles funestus*, a malaria mosquito endemic to sub-Saharan Africa, contains an impressive number of paracentric polymorphic inversions, which are unevenly distributed among chromosomes and provide an excellent framework for investigating the genomic impacts of chromosomal rearrangements. Here we present results of a fine-scale analysis of genetic variation within the genome of two weakly differentiated populations of *Anopheles funestus* inhabiting contrasting moisture conditions in Cameroon. Using population genomic analyses, we found that genetic divergence between the two populations is centered on regions of the genome corresponding to three inversions, which are characterized by high values of *F*_ST_, absolute sequence divergence and fixed differences. Importantly, in contrast to the 2L chromosome arm, which is collinear, nucleotide diversity is significantly reduced along the entire length of three autosome arms bearing multiple overlapping chromosomal rearrangements. These findings support the idea that interactions between reduced recombination and natural selection within inversions contribute to sculpt nucleotide polymorphism across chromosomes in *An. funestus*.

## Introduction

Although much progress has been made in analyzing genetic variation among populations, less is known about the complex interactions between the different driving forces — including mutation, selection, recombination, gene flow, demography or incomplete lineage sorting — which shape nucleotide polymorphism across genomes (Begun & Aquadro 1992; Betancourt & Presgraves 2002; Nordborg & Tavar 2002; Comeron *et al*. 2008; Cutter & Payseur 2013; Gosset & Bierne 2013; Huber *et al*. 2014; Cruickshank & Hahn 2014). Most genetic variability is neutral or nearly so (Kimura 1983), but patterns of polymorphism within species are also influenced by several types of natural selection whose signatures are heterogeneous in nature and across the genome (Hill & Robertson 1966; Maynard Smith & Haigh 1974; Hudson & Kaplan 1988; Charlesworth *et al*. 1993; Gillespie 1997). One fundamental, unresolved question is how the adaptation of different gene pools to different fitness conditions imposed by spatially varying natural selection in heterogeneous environments (local adaptation) affects their genomic variation (Williams 1966; Kawecki & Dieter 2004; Savolainen *et al*. 2013).

Population genetics theory and empirical models suggest that candidate genomic regions, which contain genes underlying responses to spatially heterogeneous selection pressures exhibit high divergence among populations, skewed polymorphism or an extended correlation among alleles from different loci (linkage disequilibrium) (Lewontin & Krakauer 1973; Schlötterer 2002; Beaumont 2005; Nielsen 2005; Storz 2005; Nosil *et al*. 2009). Because migration, gene flow and recombination among diverging populations can break down locally adapted allelic combinations, genetic targets of local adaptation are often protected within genomic regions of reduced recombination – including chromosomal inversions (Noor *et al*. 2001; Rieseberg 2001; Ortiz-Barrientos *et al*. 2002; Butlin 2005; Kirkpatrick & Barton 2006; Joron *et al*. 2011; Roesti *et al*. 2012, 2013; Yeaman 2013; Nishikawa *et al*. 2015).

Structural rearrangements known as chromosomal inversions, which occur when a piece of DNA within a single chromosome breaks and rotates 180° before being reinserted in the reversed orientation (Sturtevant 1921), are widespread in plants and animals (Dobzhansky 1970). A large body of literature supports a major implication of paracentric inversions (which do not encompass the centromere) in evolutionary adaptation in a wide variety of species (see Krimbas & Powell 1992; Powell 1997; Hoffmann *et al*. 2004; Hoffmann & Rieseberg 2008; Kirkpatrick 2010 for a review). This includes many examples of polymorphic inversions whose frequencies are correlated with environmental variables and temporal changes consistent with natural selection in dipteran species (Dobzhansky 1943; Mettler *et al*. 1977; Coluzzi *et al*. 1979; Knibb 1982; Krimbas & Powell 1992; De Jong & Bochdanovits 2003; Schaeffer *et al*. 2003; Anderson *et al*. 2005; Schaeffer 2008). The adaptive potential of some rearrangements is also highlighted by experimental evolution and phenotypic studies showing a remarkable association between the inversion and several fitness-related traits (Wright & Dobzhansky 1946; Dobzhansky 1948; Rako *et al*. 2006; Hoffmann & Weeks 2007; Kennington *et al*. 2007; Lowry & Willis 2010; Lee *et al*. 2011; Fouet *et al*. 2012; Kapun *et al*. 2014, 2016).

Another important property of inversions concerns the significant role they play in genome evolution as a whole. Recombination is strongly reduced between two paired chromosomes that differ by an inversion through several mechanisms, which impede crossing over (reviewed in Roberts 1976). Local recombination rates affect a myriad of processes including selection, gene conversion, diversity and divergence throughout the genome (Begun & Aquadro 1992; Kliman & Hey 1993; Andolfatto *et al*. 2001; Nordborg *et al*. 2005; Haddrill *et al*. 2007; Kulathinal *et al*. 2008; Comeron *et al*. 2012; Nachman & Payseur 2012; Roesti *et al*. 2012, 2013; Campos *et al*. 2014; Ortiz-barrientos *et al*. 2016). Since the pioneering work of Begun and Aquadro (1992), which demonstrated a positive correlation between recombination rate and nucleotide diversity within the genome of *Drosophila melanogaster*, multiple studies have provided empirical evidence that reduced crossing over alters genetic variation along chromosomes in a wide variety of organisms (Hellmann *et al*. 2003, 2005; Tenaillon *et al*. 2004; Begun *et al*. 2007; Kulathinal *et al*. 2008, 2009; Pegueroles *et al*. 2010; Corbett-Detig & Hartl 2012; McGaugh *et al*. 2012; Pool *et al*. 2012). Consequently, inversions, which are coldspots of recombination, have the potential to modulate patterns of genetic variation across relatively significant regions of the genome. More specifically, the presence of one or several rearrangements may reduce crossing over and diversity along an entire chromosome or chromosome arm, which in turn evolves differently from the rest of the genome. This scenario is best illustrated by the nonrecombining part of the Y chromosome in mammals, which is maintained by multiple inversions that have suppressed recombination over the entire length of the chromosome (Lahn & Page 1999). The potential of inversions to reduce recombination and diversity in large regions of the genome has long been exploited to engineer chromosomes or chromosomal arms with multiple inversions that block recombination, allowing lethal or sterile mutations to be stably maintained in a heterozygous state (balancer chromosomes) (Muller 1918; Sturtevant 1921; Hentges & Justice 2004; Edgley *et al*. 2006). Despite these clear examples, our understanding of how inversions affect polymorphism over the entire length of a chromosome remains limited in part due to the lack of empirical support across taxa. Targeted studies based on fine-scale examinations of genomic variation in species with well-characterized patterns of naturally occurring inversions are important for assessing both the genomic extent and the ubiquity of diversity reduction associated with chromosomal rearrangements.

An excellent opportunity to investigate the role of inversions in genome evolution exists in *An. funestus*, the second most important vector of *Plasmodium* parasites in the Afrotropical region behind the best known species, *An. gambiae* (Gillies & De Meillon 1968; Gillies & Coetzee 1987; Sinka *et al*. 2010; Coetzee & Koekemoer 2013). The two taxa share several characteristics including a near continental distribution, a marked preference for human hosts and intradomicillary host-seeking and resting behavior, which reflect their efficiency as vector of malaria parasites (Dia *et al*. 2013; Lanzaro & Lee 2013). Numerous studies have suggested that ongoing adaptive divergence contributes in part to the significant environmental flexibility, which underlies the widespread distribution of *An. funestus* populations. This species has been described as an amalgam of at least two relatively differentiated ecotypes whose ecology, phenotypic divergence, distribution range and role in malaria transmission remain obscure (Green & Hunt 1980; Costantini *et al*. 1999; Dia *et al*. 2000; Kamau *et al*. 2002; Boccolini *et al*. 2005; Cohuet *et al*. 2005; Michel *et al*. 2005; Ayala *et al*. 2011; Barnes *et al*. 2017). A very high number of paracentric polymorphic chromosomal inversions have been identified in polytene chromosomes of *An. funestus* via cytogenetic studies (Green & Hunt 1980; Sharakhov *et al*. 2004). Some of these rearrangements are spread along stable geographic clines and exhibit a significant deficit in heterokaryotypes, suggesting that they may play a role in environmental adaptation (Costantini *et al*. 1999; Dia *et al*. 2000; Kamau *et al*. 2002; Boccolini *et al*. 2005; Ayala *et al*. 2011). Importantly, dozens of large rearrangements have been detected on all autosomes except for 2L (Sharakhov *et al*. 2004). This uneven distribution of inversions between chromosome arms provides an ideal framework for testing hypotheses about the impacts of rearrangements at the chromosome level via cross-chromosomal comparisons.

In this study, we have used Restriction site Associated DNA Sequencing (RAD-Seq) (Baird *et al*. 2008) to address the effects of inversions on the genetic architecture of selection and diversity across the genome of *An. funestus* populations collected from Cameroon. Although the genome-wide level of differentiation is weak, signals of divergence are apparent within three large chromosomal rearrangements whose frequencies vary along moisture gradients in this region (3R*a*, 3R*b* and 2R*h*). Genome scans show that, in contrast to the collinear 2L arm, the proportion of polymorphic sites is drastically reduced along the entire length of three autosomal arms bearing multiple polymorphic inversions. These findings suggest that the evolution of chromosomes is impacted by the combined effects of suppressed recombination and selection within paracentric polymorphic inversions in *An. funestus*.

## Materials and methods

### Mosquito samples

Populations of *An. funestus* that occur in Cameroon have been assimilated to two weakly differentiated ecotypes distributed along a moisture gradient spanning the whole country (Costantini *et al*. 1999; Cohuet *et al*. 2005; Ayala *et al*. 2011). We sequenced 132 mosquitoes collected from two locations separated by ~500km, representative of the savannah and the forest ecogeographic domains (Fig. 1A and Table S1). In Mfou, a small city of the forest region, *An. funestus* mosquitoes breed in an artificial lake and maintain abundant populations throughout the year. In Tibati, which lies within the forest-savannah transition zone, several artificial lakes also provide abundant breeding sites for dense and perennial populations of *An. funestus*. Between August and November 2013, we used several sampling methods (Service 1993) to collect *An. funestus* larvae in breeding sites and adults seeking human hosts in and around human dwellings at night, or resting indoors during daytime. (Fig. 1A and Table S1). With this diversified sampling, we aimed to maximize the chances that our sample represents a good approximation of the genetic diversity of *An. funestus* populations, as genetic differentiation within malaria vectors species sometimes overlap with subtle microgeographic and temporal segregations (e.g. Riehle *et al*. 2011). *An. funestus* belong to a large taxonomic unit comprising at least seven taxa that are identified by slight morphological variations and a diagnostic PCR based on mutations of the ribosomal DNA (Gillies & De Meillon 1968; Gillies & Coetzee 1987; Cohuet *et al*. 2003). We verified, using morphology and PCR, that all samples included in this study belonged to the nominal species and only important malaria vector of the group: *An. funestus*.

### Library preparation, sequencing and SNP identification

We extracted genomic DNA of larvae and adult specimens with the DNeasy Blood and Tissue kit (Qiagen) and the Zymo Research MinPrep kit, respectively. Double-digest Restriction-site Associated DNA (ddRAD) libraries were prepared as described in Kamdem *et al*. (2017) following a modified protocol of Peterson *et al*. (2012) and single-end sequenced to 100 base reads on Illumina HiSeq2000.

Illumina sequences were sorted according to barcode tag and filtered using the *process_radtags* program of the Stacks v 1.35 software (Catchen *et al*. 2013). Reads with ambiguous barcode, inappropriate restriction site or low sequencing quality score were removed. We checked the depth of sequencing coverage of the final data set after all filtering steps using VCFtools (Danecek *et al*. 2011). To call and genotype SNPs, we first aligned the remaining high-quality reads to the *An. funestus* reference sequence with GSNAP (Wu & Nacu 2010), with a maximum of five mismatches allowed and terminal alignments prevented. We then used Stacks to build consensus RAD loci and to identify SNPs within each locus. We set the minimum number of reads required to form a “stack” to three and allowed two mismatches during catalogue creation. Following assembly and genotyping, the polymorphism data was further filtered to maximize data quality. To do this, we retained only RAD loci scored in every population and in at least 60% of individuals within each population and we randomly selected one SNP per locus for further analyses. SNP files in different formats used for downstream analyses were created with the *populations* program in Stacks, PLINK v 1.09 and PGDSpider v 2.0.8.2 (Purcell *et al*. 2007; Lischer & Excoffier 2012; Catchen *et al*. 2013).

### Population structure and genetic divergence

We examined the genetic relatedness among individuals with a Principal Component Analysis (PCA), a neighbor-joining (NJ) tree and the STUCTURE v 2.3.4 software (Pritchard *et al*. 2000) using SNPs identified in Stacks. We utilized the R package *adegenet* to implement the PCA and *ape* to compute a genotype-based Euclidian distance matrix between individuals and to infer individual-based NJ networks (Paradis *et al*. 2004; Jombart 2008; R Development Core Team 2016). In STRUCTURE, we ran five replicates of 1 to 10 assumed clusters (k). Each run consisted of 200000 iterations, and the first 50000 iterations were discarded as burn-in. CLUMPP v1.1.2 (Jakobsson & Rosenberg 2007) was used to aggregate results across multiple STRUCTURE runs and the clustering results were visualized graphically using DISTRUCT v1.1 (Rosenberg 2004). To find the optimal number of genetically distinct clusters, we used both the Discriminant Analysis of Principal Component (DAPC) implemented in *adegenet* and the ad hoc statistic DeltaK of Evanno *et al*. (2005) (Evanno *et al*. 2005; Earl & VonHoldt 2012).

To assess the level of genetic differentiation between savannah and forest populations, we estimated the overall *F*_ST_ (Weir & Cockerham 1984) in Genodive v1.06 (Meirmans & Van Tienderen 2004) using a subset of 2000 randomly selected SNPs. To examine to what extent the geographic region contributes to the genetic variance among samples, we conducted a hierarchical analysis of molecular variance (AMOVA) (Excoffier *et al*. 1992) in GenoDive. We used 10000 permutations to assess the statistical significance of *F*_ST_ and AMOVA.

### Genomic targets of selection

The current *An. funestus* draft genome assembly consists of 12243 scaffolds ranging from 2334 to 3334433bp in length (Giraldo-Calderon *et al*. 2015; Neafsey *et al*. 2015). Therefore, to perform genome scans and inspect footprints of selection throughout the genome, we ordered and concatenated 104 long scaffolds whose positions on the physical map have been inferred via alignment and orthology (Neafsey *et al*. 2015) and we created “pseudo-chromosomes” corresponding to the five chromosome arms of *An. funestus*. These mapped scaffolds accounted for 33% in length of the approximate size of the *An. funestus* genome (Fig. S1 and Table S2).

To delineate genomic signatures of selection, we performed an outlier analysis in order to detect genomic regions exhibiting exceptional differentiation or diversity that are putative targets of selection (Lewontin & Krakauer 1973; Storz 2005). Locus-specific *F*_ST_ values between savannah and forest populations were estimated with the *populations* program in *Stacks* and visualized graphically using the subset of SNPs located on mapped scaffolds. SNPs with *F*_ST_ values above the top 1% of the empirical distribution were considered as outliers of genetic differentiation. Loci with unusually low or high *F*_ST_ values relative to neutral expectations were also detected using the coalescence-based method FDIST2 (Beaumont & Nichols 1996) as implemented in LOSITAN (Antao *et al*. 2008). The mean neutral *F*_ST_ across all SNPs was approximated by choosing the “neutral mean *F*_ST_ option” with 99% confidence interval in LOSITAN. We ran LOSITAN with 100000 simulations and assumed a false discovery rate of 0.1 to minimize the number of false positives. *F*_ST_ values are dependent on within-population genetic diversity, which may bias estimates of the level of divergence among populations (Noor & Bennett 2009; Cruickshank & Hahn 2014). To alleviate this effect in our analyses, we used two complementary statistics to assess the degree of genetic divergence among populations across the genome — the absolute sequence divergence (*d*_xy_) and the proportion of fixed differences between populations (*d*_f_). Both statistics were estimated in ngsTools using genotype likelihood without SNP calling (Fumagalli *et al*. 2014), and kernel smoothed values were visualized in non-overlapping 90-kb windows along pseudo-chromosomes in R.

To inspect genomic patterns of genetic diversity and allele frequency spectra, we calculated pairwise nucleotide diversity (θ _π_), Watterson’s estimator of segregating sites (θ_w_) and Tajima’s *D* across RAD loci located on mapped scaffolds in ANGSD v 0.612 (Korneliussen *et al*. 2014). This program derives diversity and allele frequency spectrum statistics using genotype likelihoods without SNP calling, thereby alleviating some of the uncertainties and biases associated with SNP calling in low coverage Next Generation Sequencing (Korneliussen *et al*. 2013). To gain a genome-wide view and identify genomic regions of exceptional diversity and/or skewed allele frequency spectra, average values of θ_π_, θ_w_ and Tajima’s *D* were determined in non-overlapping 90-kb windows and plotted along pseudo-chromosomes.

## Results

### Genetic differentiation within *An. funestus*

Using alignments to the draft reference genome, we assembled 490871 unique RAD loci. We identified a total of 10687 high-quality biallelic markers by randomly choosing one SNP across loci that were present in all populations and in at least 60% of individuals within every population. A NJ tree and the first three PCA axes based on these variants clearly distinguished two genetic clusters likely corresponding to the two ecotypes previously described in Cameroon and in several other countries with a diversity of genetic markers (Costantini *et al*. 1999; Dia *et al*. 2000; Kamau *et al*. 2002; Boccolini *et al*. 2005; Cohuet *et al*. 2005; Michel *et al*. 2005; Ayala *et al*. 2011; Barnes *et al*. 2017) (Fig. 1B and 1C). The method of Evanno *et al*. (2005) and DAPC confirmed the occurrence of two to three distinct gene pools reflecting the ecological divergence known between forest and savannah populations in Cameroon (Fig. 1D and 1E). However, despite this apparent geographic segregation, the overall genetic differentiation is low between the two putative subgroups (*F*_ST_ = 0.033, *p* < 0.005). Consistent with weak genetic divergence, STRUCTURE analyses revealed a single cluster of individuals with admixed ancestry at k = 2 (Fig. 1F). The moderate genetic differentiation between savannah and forest populations is also illustrated by the results of an AMOVA, which show that the greatest proportion of the genetic variance (98.2%, *p* < 0.005) among our samples is explained by within-individual variations. The geographic origin of individuals accounts for less than 2% of the genetic variance. Overall, the genetic differentiation of *An. funestus* in Cameroon suggests a low level of genomic divergence, consistent with extensive gene flow and/or recent split between the two putative ecotypes.

**Figure 1:**
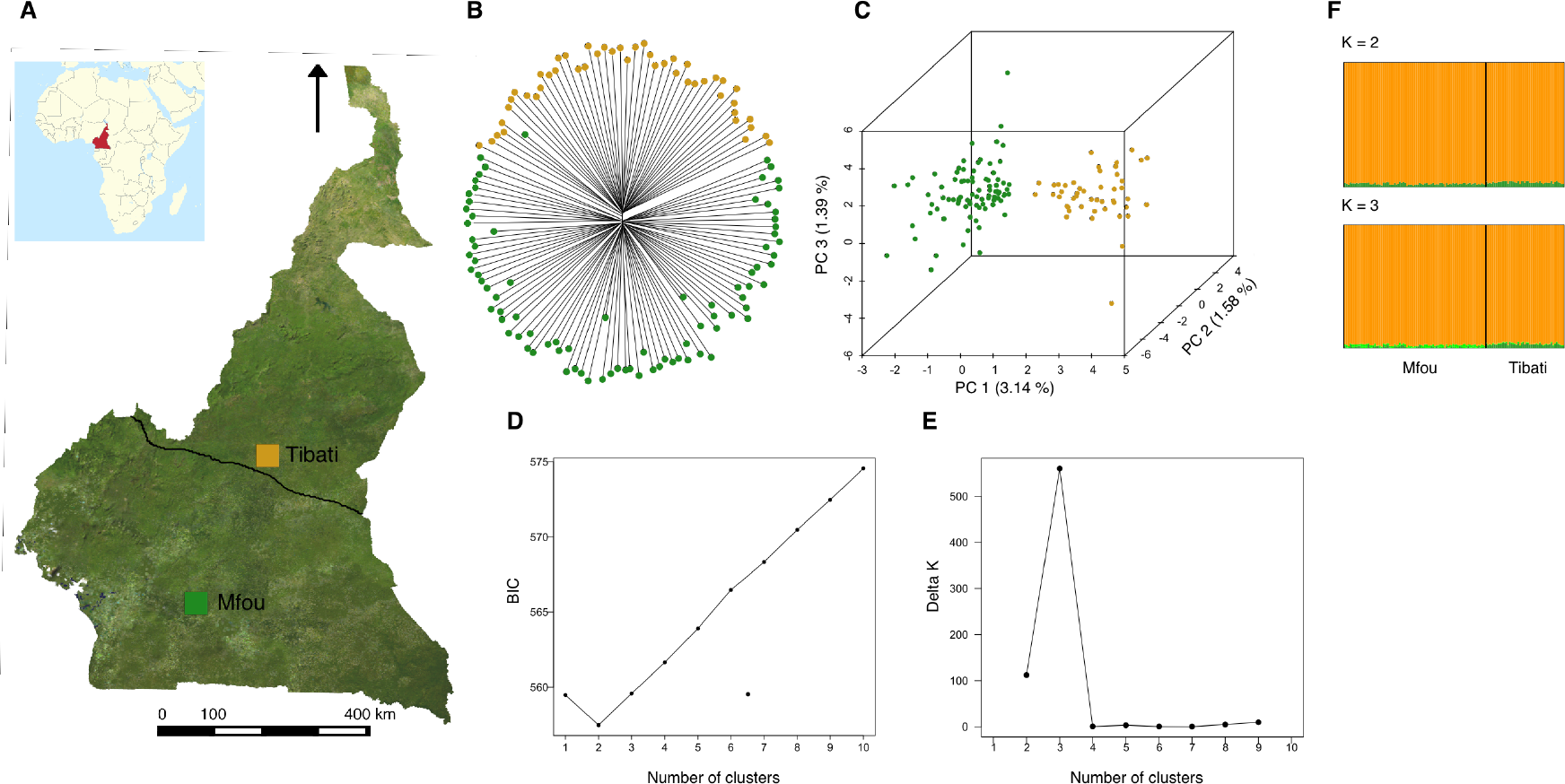
Geographic origin and genetic relatedness of *An. funestus* populations in Cameroon. (A) Map showing Mfou and Tibati, the two locations where mosquitoes were collected. A delimitation of the approximate distribution ranges of the two ecotypes that occur in Cameroon is shown (continuous line in the middle of the map). (B) and (C) Population genetic structure as revealed respectively by a neighbor-joining tree and a PCA. The percentage of variance explained is indicated on each PCA axis. (D) and (E) Confirmation of the presence of two *An. funestus* clusters with DAPC and the delta k method of Evanno *et al*. (2005). The lowest Bayesian Information Criterion (BIC) and cross-validation error and the highest delta k indicate the most probable number of clusters (2-3). (F) Bayesian clustering in STRUCTURE.

### Divergence within chromosomal inversions

We scanned the genome of savannah and forest populations to identify the few regions of the genome that diverge within a largely homogeneous background. We found that, in our panel of 10687 SNPs, highly differentiated loci are non-randomly distributed, the highest *F*_ST_ values being clustered in genomic regions bearing known polymorphic chromosomal inversions that have been previously verified in specimen collected from this area (Cohuet *et al*. 2005; Ayala *et al*. 2011) (Fig. 2 and Fig. S1). We identified a total of 107 outliers that fell above the 99^th^ percentile of the empirical distribution of *F*_ST_, including 31 SNPs that were successfully mapped to concatenated scaffolds and were confirmed as statistical outliers in LOSITAN. *F*_ST_ outliers were located exclusively on two chromosome arms containing nearly 20 polymorphic inversions (3R and 2R) (Fig. 2 and Fig. S1) (Sharakhov *et al*. 2004). Moreover, in contrast to 2L, 3L and X, which have no discriminatory power, SNPs that mapped to 3R or 2R reproduce the segregation observed between the savannah and the forest at the genome level (Fig. 3). The number of SNPs with *F*_ST_ values above the 1% threshold also varies between the 2R and 3R (9 against 22) implying that mutations or structural variants of the 3R arm are more strongly correlated with the genetic differentiation between the two populations. Overall, the genomic distribution of *F*_ST_ outliers suggests that divergent genomic regions between ecotypes are located within polymorphic inversion on two chromosome arms.

**Figure 2:**
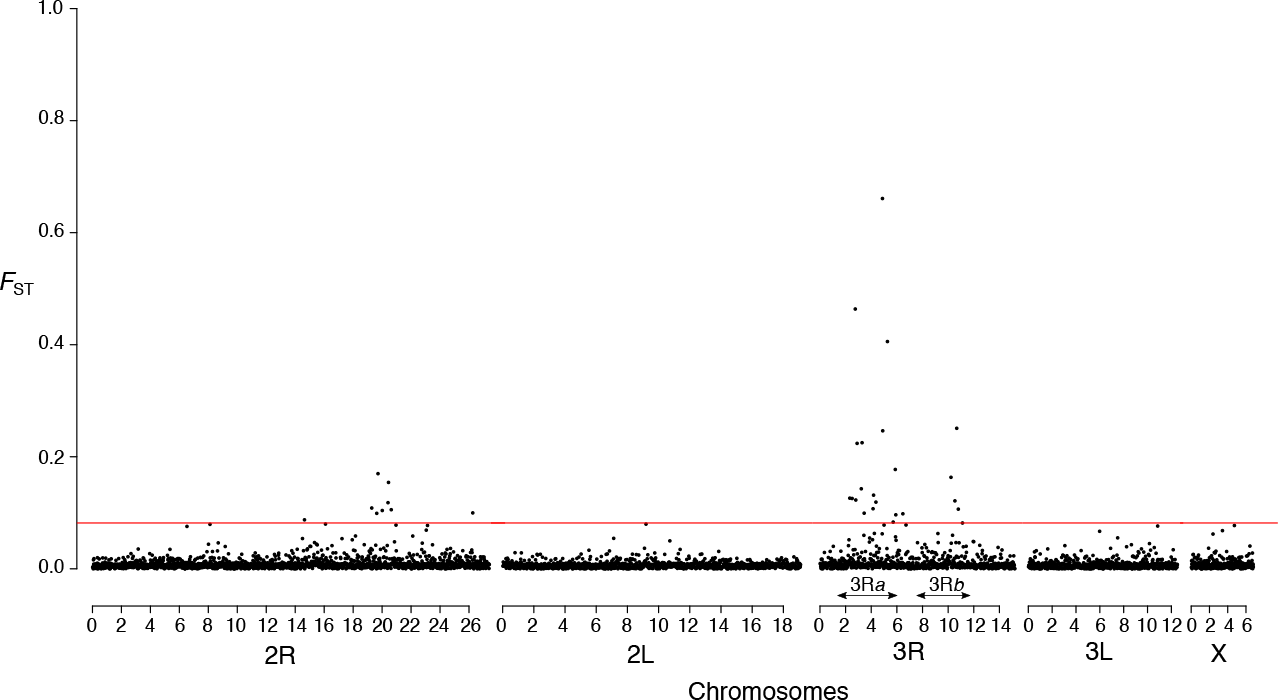
Estimates of pairwise population differentiation (*F*_ST_) based on SNPs ordered by position along pseudo-chromosomes representing the five chromosome arms of *An. funestus*. *F*_ST_ values on top of the red line are above the 99^th^ percent of the empirical distribution. Arrows indicate the genomic coordinates of the 3R*a* and 3R*b* inversions.

To further explore the role of inversions in genetic divergence, we examined the population structure separately for three large chromosomal rearrangements present on 3R (3R*a*, 3R*b* and 3R*d*) and for the 2R*h* that occurs on 2R. Although several inversions coexist and overlap along 2R, six out of nine *F*_ST_ outliers identified on this chromosome mapped to scaffolds specific to portions of the 2R*h* inversion, making it an interesting candidate locus. SNPs located within the 3R*a* assign more that 90% of individuals to their respective ecotypes, and the 3R*b* also separates a significant proportion of individuals between both sampling sites (Fig. 3). In addition, consistent with the genomic pattern of differentiation, which indicates that 83% of *F*_ST_ outliers map to these two inversions (Fig. 2), the genome-wide distribution of *d*_xy_ and fixed differences shows clear peaks within these two large inversions (Fig. 4). The population structure of 2R*h* is comparable with that of 3R*b*, suggesting that this inversion also contributes to the savannah-forest segregation of *An. funestus* populations in Cameroon. Conversely, SNPs identified outside the 3R*a* and 3R*b*, or within the 3R*d* have no discriminatory power (Fig. 3). In summary, the population structure of inversions on 2R and 3R reflects the genomic distribution of differentiated loci, which suggests that divergence centered on three chromosomal rearrangements (3R*a*, 3R*b* and 2R*h*) maintains some level of genomic integrity despite low genome-wide differentiation between ecotypes of *An. funestus* in this geographic area. This divergence translates into *F*_ST_ values, which increase from 0.033 at the genome level to 0.053, 0.08 and 0.22 in 2R*h*, 3R*b* and 3R*a*, respectively. Similarly, the proportion of the genetic variance explained by the geographic origin of samples rises from 1.8% for the genome-wide SNPs to 3.2%, 4.7% and 12.9% when only variants present within the 2R*h*, 3R*b* and 3R*a* rearrangements, respectively, are included.

**Figure 3:**
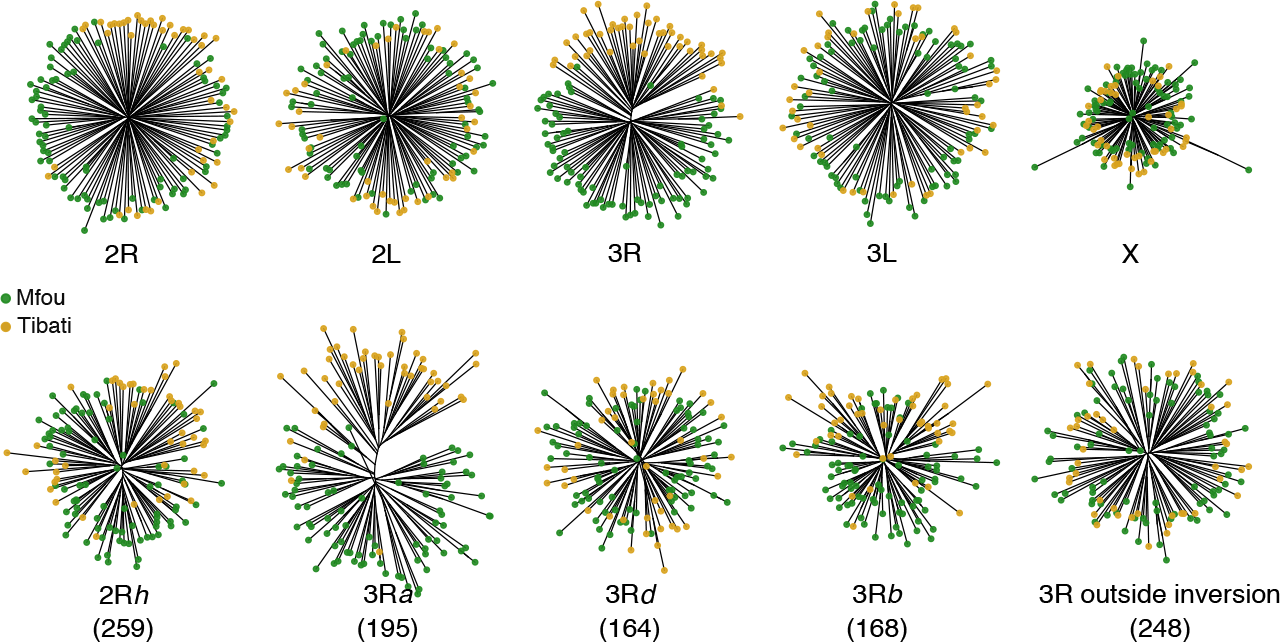
Population genetic structure revealed by each chromosome arm and by four polymorphic chromosomal inversions (2R*h*, 3R*a*, 3R*b* and 3R*d*). The number of SNPs is indicated in parenthesis.

**Figure 4:**
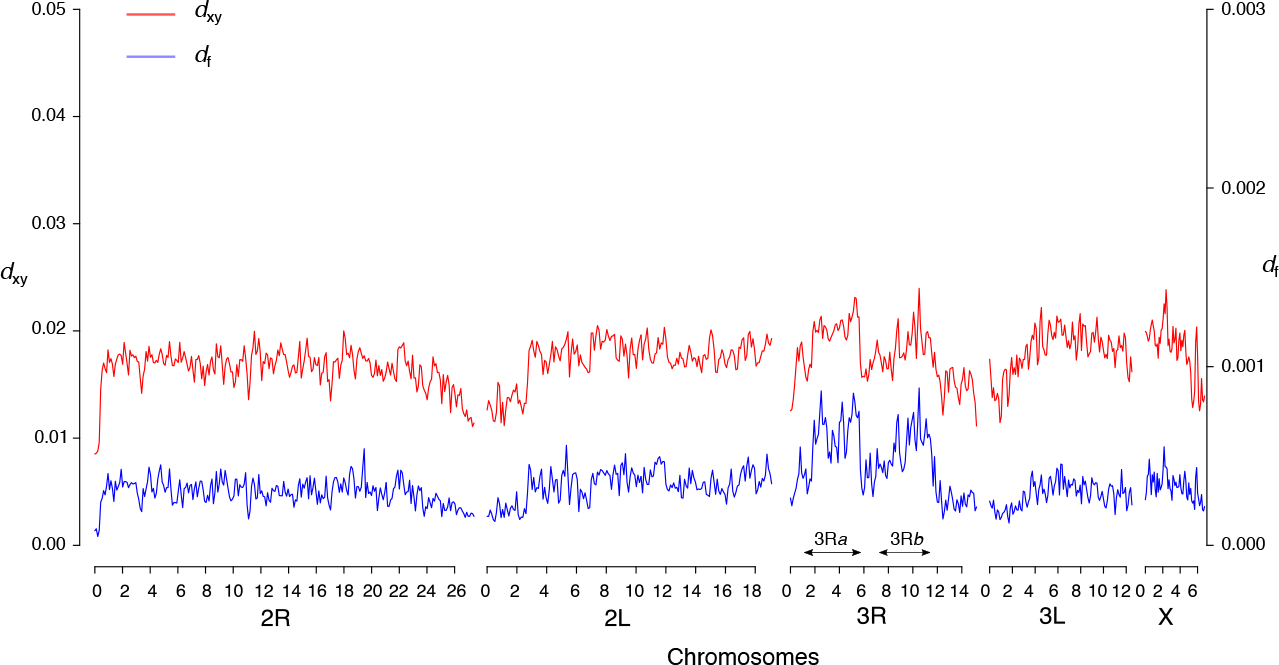
Genome-wide distribution of *d*_xy_ and fixed differences (*d*_f_) across 90-kb non-overlapping windows along the five pseudo-chromosomes in Mfou and Tibati populations. Strong sequence divergence within the 3R*a* and 3R*b* inversions is characterized by the presence of peaks of *d*_xy_ and *d*_f_ at these loci.

### Chromosome arm-specific diversity associated with inversions

We investigated patterns of polymorphism across the genome using scans based on estimates of nucleotide diversity (θ_π_ and θ_w_) and Tajima’s *D* (Fig. 5). The proportion of polymorphic sites (θ_w_) varies substantially between chromosome arms, the highest values being found on the only collinear chromosome arm (2L) (Fig. 5 and Fig. 6). Precisely, the average value of θ_w_ differs significantly between 2L and each of the three other autosomes (Wilcoxon rank sum test, *p* < 0.001) (Fig. 6). θ_w_ is reduced by 34.1% on 3R, 24.0% on 2R and 13.2% on 3L relative to the 2L arm, regardless of the sampling location. Estimates of the number of polymorphic sites along each of the five chromosome arms are also significantly different between Mfou (forest) and Tibati (savannah) populations (Wilcoxon rank sum test, *p* < 0.001). Additionally, the drop of the average θ_w_ on inverted autosomes relative to 2L is 34.7% on 3R, 29.2% on 2R and 13.2% on 3L in Mfou compared with 33.5% on 3R, 18.8% on 2R and 13.2% on 3L in Tibati (Fig. 5 and Table 1). A different demographic history between ecotypes or diverse other factors including the variation in sample size between the two locations can account for this difference in the level of polymorphism between populations. Nevertheless, since most of the polymorphic chromosomal inversions (notably 3R*a* and 3R*b*) are fixed in forest populations and fluctuate in Tibati (Cohuet *et al*. 2005; Ayala *et al*. 2011), it is likely that variations in karyotype frequencies of inversions contribute at least in part to the stronger reduction of nucleotide diversity on inverted autosomes observed in Mfou compared with Tibati. The X chromosome bears no known chromosomal rearrangement, but likely due to its particular divergence and its distorted effective population size, its average θ_w_ is also weak compared with 2L (21.1% reduction relative to 2L) regardless of the sampling site. A significant reduction in pairwise nucleotide diversity (θ_π_) relative to the 2L is found on chromosomes 3R, X and 2R to a lesser extent, but there are no clear patterns among chromosomes and sampling locations as is the case with θ_w_ (Fig. 5, Fig. 6 and Table 1).

**Figure 5:**
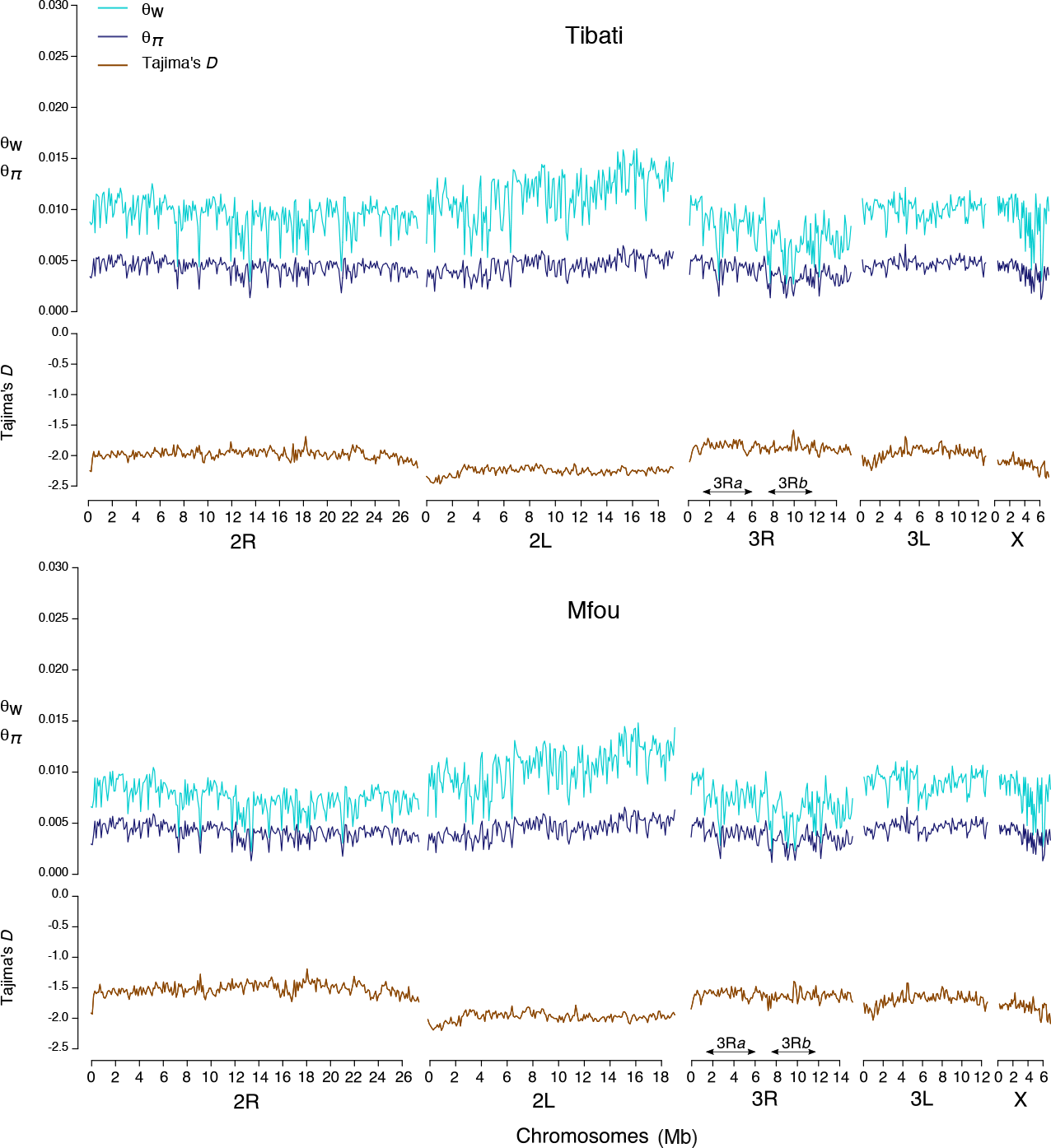
Estimates of nucleotide diversity (θ_π_ and θ_w_) and allele frequency spectrum (Tajima’s *D*) across 90-kb non-overlapping windows illustrating the uneven distribution of genetic diversity along the five chromosome arms. The collinear autosome (2L) is the most polymorphic in both Mfou and Tibati populations. In contrast, autosomes bearing multiple inversions exhibit a significant reduction in the amount of polymorphic sites.

**Figure 6:**
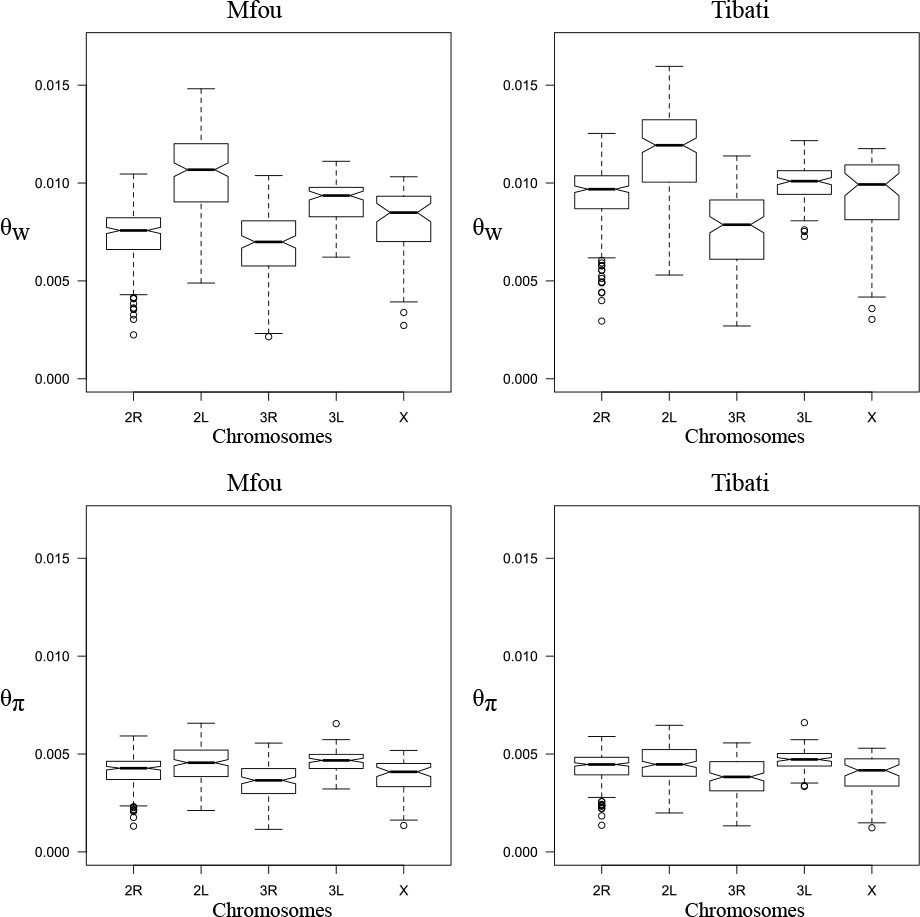
Box plot depicting the distribution of θ_π_ and θ_w_ among chromosomes in Mfou and Tibati.

**Table 1:** Reduction in nucleotide diversity relative to the 2L chromosome arm (%).

|  |  | **2R** | **3L** | **3R** | **X** |
| --- | --- | --- | --- | --- | --- |
| Mfou | $\theta_w$ | 29.22 * | 13.24 * | 34.69 * | 23.35 * |
| | $\theta_\pi$ | 7.92 * | -3.23 | 19.23 * | 13.92 * |
| Tibati | $\theta_w$ | 18.79 * | 13.24 * | 33.54 * | 18.94 * |
| | $\theta_\pi$ | 3.23 | -4.84 | 15.61 * | 11.86 * |

\*Statistically significant (Wilcoxon rank sum test, $p < 0.001$ )

Local recombination rates are strongly correlated with genetic diversity and the effective population size (Begun & Aquadro 1992). Therefore, it is possible and parsimonious that the strong variation in polymorphism observed among autosomes in *An. funestus* is due to the presence of multiple overlapping inversions that are under selection in different geographic contexts throughout the species’ range. These inversions reduce meiotic crossover and genetic variation along the 2R, 3R and 3L autosomes, which contain multiple rearrangements. For instance, three inversions (3R*a*, 3R*b* and 3R*d*) account for at least 75% of the length of the 3R arm (Fig. S1). Similarly, nearly 80% of the length of 2R and 3L is affected by chromosomal rearrangements. The idea that the combined effects of low recombination and selection within multiple inversions contributes to reduce polymorphism throughout the three inverted autosomal arms is supported by the pattern of diversity observed on the collinear chromosome arm 2L, which acts as a negative control. Conceivably, polymorphism is highest on this chromosome arm due to the absence of inversions, which preserves more substantial levels of recombination.

The genome-wide diversity falls in the range described in other highly polymorphic *Anopheles* populations with RADseq markers (θ_w_ = 0.0083 and θ_π_ = 0.0042 in Mfou; θ_w_ = 0.0097 and θ_π_ = 0.0043 in Tibati) (O’Loughlin *et al*. 2014; Fouet *et al*. 2017; Kamdem *et al*. 2017). The range of values of Tajima’s *D* (from - 2.47 to -1.58 among Tibati samples and from -2.20 to -1.19 in Mfou) was shifted towards negative values suggestive of recent population expansion leading to an accumulation of low-frequency variants, although the effects of population structure, selective sweeps, background selection or changes in sample size can contribute to skewed site frequency spectra across the genome (Donnelly *et al*. 2001; Thornton 2005; Gattepaille *et al*. 2013) (Fig. 5). Unsurprisingly, Tajima’s *D* is more negative on the 2L chromosome, which exhibits the greatest difference between the number of polymorphic sites and the pairwise nucleotide diversity in both ecotypes.

## Discussion

### Recombination, inversions and genetic divergence

Our analysis based on more than 10000 genome-wide SNPs revealed that *An. funestus* populations collected from two different moisture conditions in Cameroon are very weakly differentiated. These results corroborate two previous studies based on microsatellites and polymorphic chromosomal inversions, which have found similar levels of divergence between *An. funestus* populations in this region (Cohuet *et al*. 2005; Ayala *et al*. 2011). The population genetic structure of this mosquito throughout Africa remains largely unresolved, but based on many previous studies in different parts of the continent, it has been proposed that at least two ecotypes segregate within this species (Green & Hunt 1980; Costantini *et al*. 1999; Dia *et al*. 2000; Kamau *et al*. 2002; Boccolini *et al*. 2005; Cohuet *et al*. 2005; Michel *et al*. 2005; Ayala *et al*. 2011; Barnes *et al*. 2017). These studies have also indicated that polymorphic chromosomal inversions play a key role in ecological divergence in *An. funestus* (Green & Hunt 1980; Costantini *et al*. 1999; Dia *et al*. 2000; Cohuet *et al*. 2005; Ayala *et al*. 2011). The 3R*a* and 3R*b* inversions in particular have long been suspected to be strongly associated with ecological divergence due to the clinal distribution of alternative karyotypes and the significant deficit in heterokaryotypes observed in several countries (Costantini *et al*. 1999; Dia *et al*. 2000; Ayala *et al*. 2011). Here we provide the first genomic evidence that genetic divergence within this species is limited to a few loci bearing large chromosomal rearrangements. Our conclusions come with some caveats due to the potential limitations of the sampling scheme, the reference genome and the RAD sequencing approach we used. For instance, it is likely that some genomic signatures of selection are not detected because of the limited genomic coverage of RAD tags and pseudo-chromosomes (Arnold *et al*. 2013; Tiffin & Ross-Ibarra 2014). Bearing in mind these caveats, the clustering of genetic differentiation within the 3R*a*, 3R*b* and 2R*h* inversions appears to be consistent with the hypothesis that these rearrangements are the major genetic targets of divergent selection in Cameroon.

The phenotypic, behavioral or ecological variations associated with inversions in *An. funestus* remain unknown. In principle, the segregation between the arid zone and the humid forest may potentially involve fitness traits and phenotypes contributing to thermal tolerance. Indeed, examples of inversions whose alternative karyotypes provide selective advantages in different moisture conditions are well known in dipteran species (e.g. the 2L*a* inversion and to a lesser extent the 2R*b* in *An. gambiae*) (Coluzzi *et al*. 1979; Gray *et al*. 2009; Lee *et al*. 2009; Fouet *et al*. 2012; Cheng *et al*. 2012). Comparative cytogenetic studies also showed a high proportion of conserved genes between the 3R*b* in *An. funestus* and the 2L*a* in *An. gambiae* and between the 2R*h* in *An. funestus* and the 2R*b* in *An. gambiae* (Sharakhova *et al*. 2011). Although this hypothesis awaits thorough investigation, the two pairs of inversions may have inherited similar phenotypes via a common ancestor or via convergent evolution between *An. funestus* and *An. gambiae*.

Theoretical models and empirical data support the idea that, at early stages of divergence as in the case of *An. funestus* ecotypes, genetic differentiation is restricted to a few genomic regions because the effects of natural selection at this step are very localized (Feder *et al*. 2012; Nosil & Feder 2012; Andrew & Rieseberg 2013; Seehausen *et al*. 2014). Also, selection at a locus affects the level of genetic differentiation among populations and reciprocally the degree of subdivision within a population impacts patterns of variation at selected loci (Lewontin & Krakauer 1973; Charlesworth *et al*. 1997; Slatkin & Wiehe 1998; Majewski & Cohan 1999; Kim & Maruki 2011; Schneider & Kim 2013). In fact, it has been proposed that selection modulates levels of divergence locally and globally across the genome through two types of genetic hitchhiking — divergence hitchhiking and genomic hitchhiking (Feder *et al*. 2012). Divergent hitchhiking (the spread of a mutation and linked neutral sequences within a population) (Maynard Smith & Haigh 1974) causes the diffusion of favorable alleles at loci important for local adaptation and ecological differentiation. Limited gene flow that occurs between divergent lineages can naturally lead to genomic hitchhiking, which reflects the global accumulation of differences at the genome level through different selective or demographic mechanisms (Feder *et al*. 2012). In *An. funestus*, it is plausible that divergent hitchhiking within the three important inversions, 3R*a*, 3R*b* and 2R*h*, is so far strongly counterbalanced by extensive gene flow and the weak overall number of selected loci across the genome, leading to minimal genomic divergence among ecotypes.

Although some controversy persists (see McGaugh *et al*. 2012), the idea that strongly differentiated genomic regions between species or between populations within the same species accumulate disproportionately in regions of low genetic recombination including chromosome centers and chromosomal rearrangements has been strongly supported in many species notably humans, *Drosophila* flies, stickleback fish and maize (Hellmann *et al*. 2003, 2005; Tenaillon *et al*. 2004; Kulathinal *et al*. 2008; Cai *et al*. 2009; Keinan & Reich 2010; Nachman & Payseur 2012; McGaugh & Noor 2012; Roesti *et al*. 2012, 2013). This coincidence can be explained by a mechanical effect of recombination, which reduces genetic polymorphism within populations and thereby generates sequence divergence between populations as a simple byproduct of diversity reduction (Begun & Aquadro 1992; Roesti *et al*. 2012). Concordance between limited rate of crossing over and increased divergence may also be directly correlated with more pronounced effects of divergent selection in genomic regions in which recombination is less frequent (Charlesworth *et al*. 1997; Charlesworth 1998; Nachman 2002; Cutter & Payseur 2013; Cruickshank & Hahn 2014). Within inversions in particular, reduced recombination enhances divergent selection acting on locally adapted gene complexes, which co-segregate with or without epistatic interactions (Dobzhansky 1950; Kirkpatrick & Barton 2006).

### Recombination, inversions and genetic polymorphism

We have found chromosome arm specific patterns of polymorphism in *An. funestus* characterized by a significant difference between collinear and inverted chromosomes. We first conducted several tests to insure that uncertainties associated with our sequencing and analytical approach cannot account for the observed disparities in genetic polymorphism between chromosomes. We compared the distribution and the length of mapped scaffolds and confirmed that each chromosome arm is represented by a substantial number of long scaffolds (Table S2 and Fig. S1). The length of pseudo-chromosomes is also proportional to the length of chromosome arms, ranging from 6.8 Mb on X (the smallest) to 27.4 Mb on 2R (the largest). Therefore, we can reasonably rule out the possibility that the chromosome-bias diversity is due to a systematic error associated with our reference sequence. Additionally, mapped scaffolds are evenly distributed along the length of chromosomes, suggesting that such important variations in the amount of polymorphism between chromosome arms are not due to centromere- and/or telomere-proximal effects (Fig. S1) (Aguade *et al*. 1989; Stephan & Langley 1989). Finally, we noted that the mean depth of sequencing coverage per mapped scaffolds was consistent across the five chromosome arms (Fig. S2), implying that chromosome-specific diversity is not simply a covariate of sequencing biases.

As estimates of the number of segregating sites (θ_w_) also known as the population-scaled mutation rate (Watterson 1975) are the most drastically affected by between-chromosome variations observed in *An. funestus*, the stark contrast in the amount of polymorphism among inverted and collinear autosomes may in theory be due to lower mutation rates on inverted chromosomes. However, as shown in other insect species, such a variability in mutation rates between chromosomes or between large segments of the genome is unlikely (Begun *et al*. 2007; Keightley *et al*. 2015). Instead, the occurrence of chromosome-specific patterns of diversity in *An. funestus* is consistent with the presence of weakly recombining autosomes bearing multiple overlapping chromosomal rearrangements. These same causes produce similar effects along the non-recombining portion of the Y chromosome in mammals (Lahn & Page 1999) or along balancer chromosomes described in *Drosophila*, *Caenorhabditis* and rodent species (Muller 1918; Hentges & Justice 2004; Edgley *et al*. 2006). Indeed, population genetic analyses across genomes of diverse taxa have found a positive correlation between the rate of recombination and genetic variation (Aguade *et al*. 1989; Stephan & Langley 1989; Begun & Aquadro 1992; Hellmann *et al*. 2003, 2005; Tenaillon *et al*. 2004; Begun *et al*. 2007; Kulathinal *et al*. 2008, 2009). This correlation translates into specific patterns of diversity that can be observed at the species, genomic or chromosomal levels. At the species level, plants and some animal species, which reproduce by self-fertilization have reduced overall genomic diversity due to low genetic recombination (Nordborg *et al*. 1996; Akhunov *et al*. 2010; Cutter & Choi 2010; Andersen *et al*. 2012; Thomas *et al*. 2015). The local effects of recombination on genetic polymorphism are also particularly evident across centromere and telomeres of chromosomes and within genomic regions bearing structural rearrangements (Aguade *et al*. 1989; Stephan & Langley 1989; Andolfatto *et al*. 2001; Corbett-Detig & Hartl 2012; Pool *et al*. 2012).

The relationship between diversity and crossing over has been ascribed to two processes: selection and mutagenesis. Recombination may influence polymorphism because of new mutations created during crossing over, which increase diversity (the mutagenic effect of recombination) (Magni & Von Borstel 1962). However, multiple studies that have analyzed fine-scale recombination in humans, yeast, *Arabidopsis*, *Anopheles*, *Drosophila*, and *Caenorhabditis* have undermined the notion that recombination, or some correlate, generally exerts mutagenic effects on genomes (Betancourt & Presgraves 2002; Stump *et al*. 2005; Spencer *et al*. 2006; Wright *et al*. 2006; Begun *et al*. 2007; Noor 2008; Denver *et al*. 2009; Cutter & Choi 2010; Mcgaugh *et al*. 2012). Alternatively, recombination may affect diversity indirectly by modulating the effects of positive or negative selection across the genome. Indeed, the hitchhiking of favorable alleles (selective sweep) or the removal of recurrent deleterious mutations (background selection) affect linked neutral sequences more strongly in low-recombination genomic regions, which gives rise to a positive correlation between diversity and recombination rate (Maynard Smith & Haigh 1974; Kaplan *et al*. 1989; Begun & Aquadro 1992; Charlesworth *et al*. 1993, 1997; Nordborg *et al*. 1996; Andolfatto 2001; Nachman 2002). Another indirect effect of recombination on genetic diversity may be driven by the Hill-Roberston interference, which occurs when two sites under weak selection are in physical linkage and as a result, selection at one site interferes with selection at another site (Hill & Robertson 1966; Felsenstein 1974). Hill-Roberston interference can also be thought of as a reduction in the effective population size (*N*_e_) caused by selection at linked loci (Comeron *et al*. 2008; Cutter & Payseur 2013; Castellano *et al*. 2016). Recombination, by alleviating interference between linked sites, alleviates this reduction in *N*_e_ leading to a positive correlation between recombination rate and levels of neutral polymorphism. Because of the relationship between recombination and polymorphism, measures of the skew in the allele frequency spectrum, such as Tajima’s *D* values (Tajima 1989), are expected to positively correlate with the rate of recombination (Braverman *et al*. 1995). The *An. funestus* genome shows a strong relationship between Tajima’s *D* values and recombination, and as expected, the chromosome 2L, whose recombination is not limited by inversions, exhibits the sharpest skew in the AFS.

## Conclusions and perspectives

We found evidence that, among the many chromosomal rearrangements identified in *An. funestus*, genomic footprints of divergence are centered on three inversions that are potential targets of selection among ecotypes in Cameroon. The other rearrangements are likely either cosmopolitan or endemic inversions under selection at various geographic extents that have yet to be resolved. Our data support the idea that interactions between recombination and selection — which amplify the effects of selective sweeps or background selection within genomic regions where recombination is blocked by multiple chromosomal inversions — account for the strong disparity in nucleotide diversity observed between autosomes in this mosquito.

To deepen our understanding of the adaptive role of inversions and their contribution to chromosome specific patterns of diversity, a complete reference genome assembly and extensive sampling across the species range are needed. These resources will make it possible to design more sensitive tests including the functional characterization of footprints of selection and their detailed signatures among individuals and populations. The presence of hallmarks of both reduced recombination and linked selection across large genomic sequences in *An. funestus* highlights the important contribution of multiple aspects of linkage in genome evolution. These aspects have significant implications for the detection of genomic signatures of adaptation in species whose genome contains multiple polymorphic chromosomal inversions.

## Acknowledgements

We thank the associate editor and six reviewers for their constructive comments. This work was supported by NIH grant 1R01AI113248 to BJW.

## Author contributions

CK, CF and BJW conceived, designed and performed the experiments. CK, CF analysed the data. CK wrote the paper.

## Data Archiving Statement

Data for this study are available at: to be completed after manuscript is accepted for publication

---

**Figure S1:**
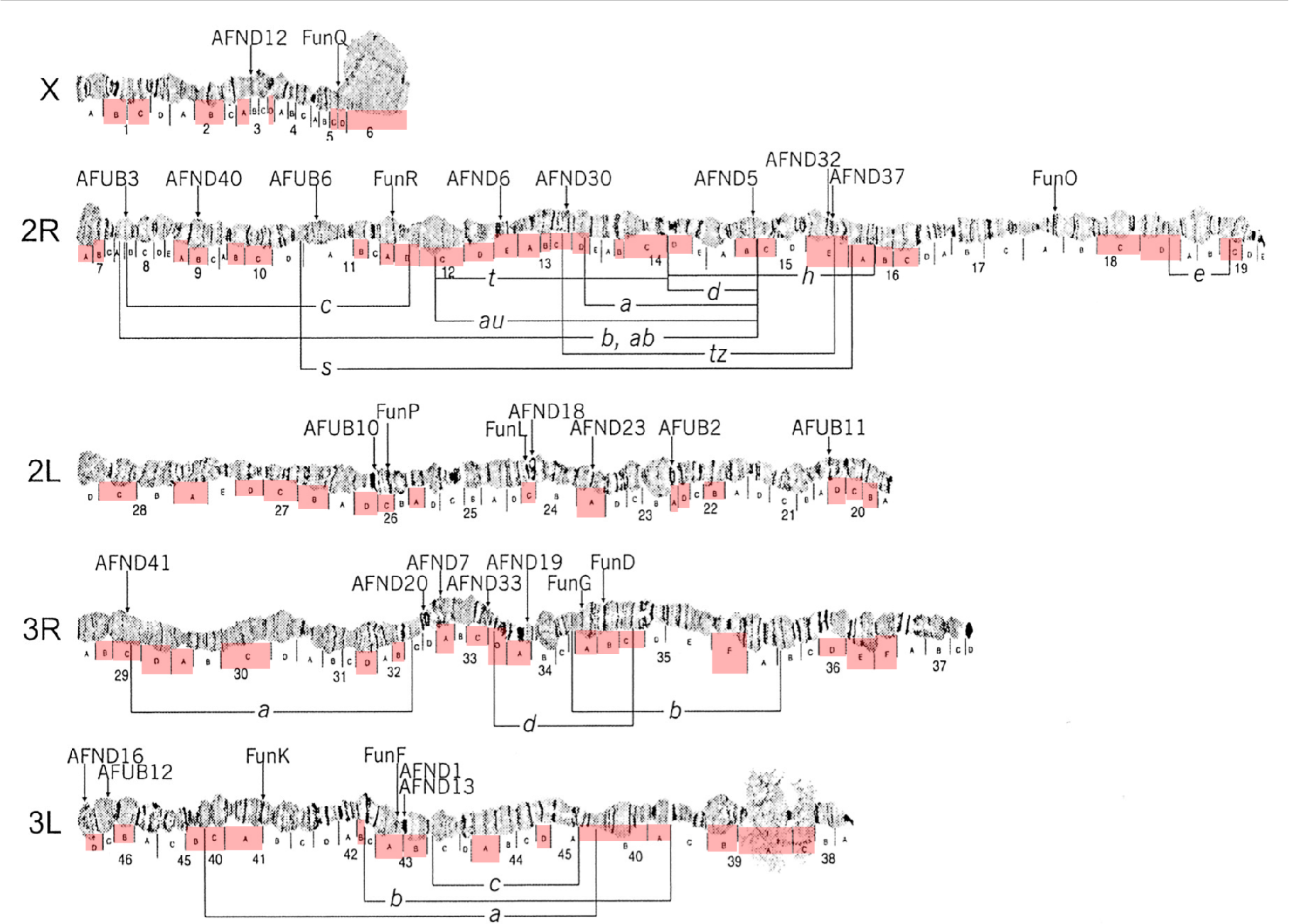
The positions of the 104 scaffolds that were placed on the *An. funestus* physical map are indicated by red rectangles. Inversions are delimitated by boxes and indicated by alphabet letters (modified from Sharakhov *et al*. 2004).

**Figure S2:**
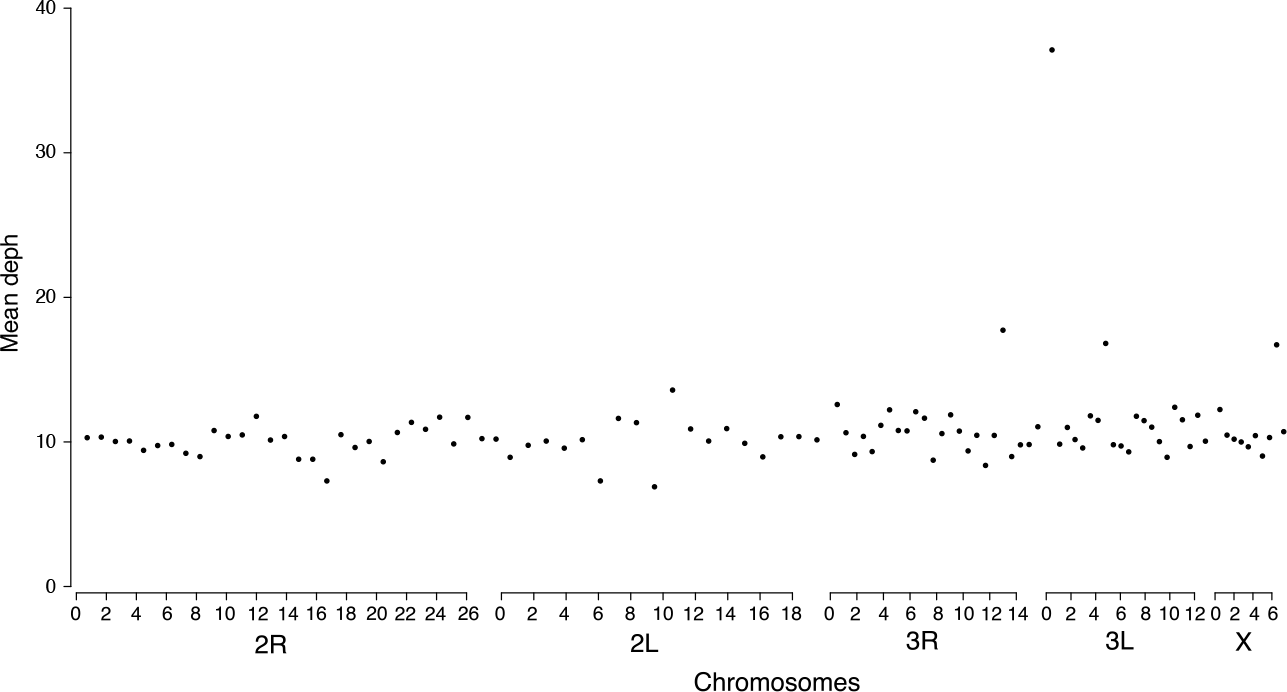
Distribution of the average coverage depth along mapped scaffolds.

**Table S1:** Description of *An. funestus* samples included in this study.

| Ecogeographic<br>region | Sampling<br>location | Geographic<br>coordinates | Sampling<br>method |  |  |  | Total | Gender |  |  |
| --- | --- | --- | --- | --- | --- | --- | --- | --- | --- | --- |
|  |  |  | HLC-OUT | HLC-IN | LC | SPRAY |  | Female | Male | NA |
| Savannah | Tibati | 6°28'08"N, 12°37'44"E | 8 | 7 | 0 | 32 | 47 | 43 | 4 | 0 |
| Forest | Mfou | 3°58'00"N, 11°56'00"E | 2 | 8 | 22 | 53 | 85 | 55 | 8 | 22 |
|  | Total |  |  |  |  |  | 132 | 98 | 12 | 22 |

HLC-OUT, human landing catches performed outdoor; HLC-IN, human landing catches performed indoor; LC, larval collection; SPRAY, spray catches; NA, not available

**Table S2:** Characteristics of the five pseudo-chromosomes of *An. funestus* used for genome scans.

| Chromosome | Number of<br>mapped<br>scaffolds | Total<br>length (bp) | Number<br>of SNPs |
| --- | --- | --- | --- |
| 2R | 30 | 27 415 570 | 1 587 |
| 2L | 18 | 19 110 819 | 987 |
| 3R | 24 | 15 197 945 | 745 |
| 3L | 22 | 12 545 136 | 652 |
| X | 10 | 6 812 527 | 296 |

## References

Aguade M, Miyashita N, Langley CH (1989) Reduced variation in the yellow-achaete-scute region in natural populations of Drosophila melanogaster. Genetics, 122, 607–15.

Akhunov ED, Akhunova AR, Anderson OD et al. (2010) Nucleotide diversity maps reveal variation in diversity among wheat genomes and chromosomes. BMC genomics, 11, 702.

Andersen EC, Gerke JP, Shapiro J a et al. (2012) Chromosome-scale selective sweeps shape Caenorhabditis elegans genomic diversity. Nature genetics, 44, 285–90.

Anderson AR, Hoffmann AA, McKechnie SW, Umina PA, Weeks AR (2005) The latitudinal cline in the In (3R) Payne inversion polymorphism has shifted in the last 20 years in Australian Drosophila melanogaster populations. Molecular Biology and Evolution, 14, 851–858.

Andolfatto P (2001) Adaptive hitchhiking effects on genome variability. Current Opinion in Genetics and Development, 11, 635–641.

Andolfatto P, Depaulis F, Navarro A (2001) Inversion polymorphisms and nucleotide variability in Drosophila. Genetical research, 77, 1–8.

Andrew RL, Rieseberg LH (2013) Divergence is focused on few genomic regions early in speciation: incipient speciation of sunflower ecotypes. Evolution; international journal of organic evolution, 67, 2468–82.

Antao T, Lopes A, Lopes RJ, Beja-Pereira A, Luikart G (2008) LOSITAN: a workbench to detect molecular adaptation based on a Fst-outlier method. BMC bioinformatics, 9, 323.

Arnold B, Corbett-Detig RB, Hartl D, Bomblies K (2013) RADseq underestimates diversity and introduces genealogical biases due to nonrandom haplotype sampling. Molecular Ecology, 22, 3179–3190.

Ayala D, Fontaine MC, Cohuet A et al. (2011) Chromosomal inversions, natural selection and adaptation in the malaria vector Anopheles funestus. Molecular biology and evolution, 28, 745–758.

Baird NA, Etter PD, Atwood TS et al. (2008) Rapid SNP Discovery and Genetic Mapping Using Sequenced RAD Markers. PLoS ONE, 3, e3376.

Barnes KG, Weedall GD, Ndula M et al. (2017) Genomic Footprints of Selective Sweeps from Metabolic Resistance to Pyrethroids in African Malaria Vectors Are Driven by Scale up of Insecticide-Based Vector Control. PLoS Genetics, 13(2): e10, 1–22.

Beaumont MA (2005) Adaptation and speciation: what can F st tell us ? Trends in Ecology & Evolution, 20, 435–440.

Beaumont MA, Nichols RA (1996) Evaluating loci for use in the genetic analysis of population structure. Proceedings of the Royal Society B: Biological Sciences, 263, 1619–1626.

Begun D, Aquadro C (1992) Levels of naturally occurring DNA polymorphism correlate with recombination rates in D. melanogaster. Nature, 356, 519–520.

Begun DJ, Holloway AK, Stevens K et al. (2007) Population genomics: whole-genome analysis of polymorphism and divergence in Drosophila simulans. PLoS biology, 5, e310.

Betancourt AJ, Presgraves DC (2002) Linkage limits the power of natural selection in Drosophila. Proceedings of the National Academy of Sciences of the United States of America, 99, 13616–20.

Boccolini D, Carrara GC, Dia I et al. (2005) Chromosomal differentiation of Anopheles funestus from Luanda and Huambo Provinces, western and central Angola. American Journal of Tropical Medicine and Hygiene, 73, 1071–1076.

Braverman JM, Hudson RR, Kaplan NL, Langley CH, Stephan W (1995) The hitchhiking effect on the site frequency spectrum of DNA polymorphisms. Genetics, 140, 783–796.

Butlin RK (2005) Recombination and speciation. Molecular Ecology, 14, 2621–2635.

Cai JJ, Macpherson JM, Sella G, Petrov DA (2009) Pervasive hitchhiking at coding and regulatory sites in humans. PLoS Genetics, 5.

Campos JL, Halligan DL, Haddrill PR, Charlesworth B (2014) The relation between recombination rate and patterns of molecular evolution and variation in drosophila melanogaster. Molecular Biology and Evolution, 31, 1010–1028.

Castellano D, Coronado-Zamora M, Campos JL, Barbadilla A, Eyre-Walker A (2016) Adaptive evolution is substantially impeded by hill-Robertson interference in drosophila. Molecular Biology and Evolution, 33, 442–455.

Catchen J, Hohenlohe P a., Bassham S, Amores A, Cresko W. (2013) Stacks: An analysis tool set for population genomics. Molecular Ecology, 22, 3124–3140.

Charlesworth B (1998) Measures of divergence between populations and the effect of forces that reduce variability. Molecular Biology and Evolution, 15, 538–543.

Charlesworth B, Morgan MT, Charlesworth D (1993) The effect of deleterious mutations on neutral molecular variation. Genetics, 134, 1289–1303.

Charlesworth B, Nordborg M, Charlesworth D (1997) The effects of local selection, balanced polymorphism and background selection on equilibrium patterns of genetic diversity in subdivided populations. Genet. Res., Camb, 70, 155–174.

Cheng C, White BJ, Kamdem C et al. (2012) Ecological genomics of Anopheles gambiae along a latitudinal cline: a population-resequencing approach. Genetics, 190, 1417–32.

Coetzee M, Koekemoer LL (2013) Molecular systematics and insecticide resistance in the major African malaria vector Anopheles funestus. Annual Review of Entomology, 58, 393–412.

Cohuet A, Dia I, Simard F et al. (2005) Gene flow between chromosomal forms of the malaria vector Anopheles funestus in Cameroon, Central Africa, and its relevance in malaria fighting. Genetics, 169, 301–311.

Cohuet A, Simard F, Toto J-C et al. (2003) Species identification within the Anopheles funestus group of malaria vectors in Cameroon and evidence for a new species. The American journal of tropical medicine and hygiene, 69, 200–205.

Coluzzi M, Sabatini A, Petrarca V, Di Deco M (1979) Chromosomal differentiation and adaptation to human environments in the Anopheles gambiae complex. Trans R Soc Trop Med Hyg, 73, 483–97.

Comeron JM, Ratnappan R, Bailin S (2012) The Many Landscapes of Recombination in Drosophila melanogaster. PLoS Genetics, 8, 33–35.

Comeron JM, Williford a, Kliman RM (2008) The Hill-Robertson effect: evolutionary consequences of weak selection and linkage in finite populations. Heredity, 100, 19–31.

Corbett-Detig RB, Hartl DL (2012) Population Genomics of Inversion Polymorphisms in Drosophila melanogaster. PLoS Genetics, 8.

Costantini C, Sagnon N, Ilboudo-Sanogo E, Coluzzi M, Boccolini D (1999) Chromosomal and bionomic heterogeneities suggest incipient speciation in Anopheles funestus from Burkina Faso. Parassitologia, 41, 595–611.

Cruickshank TE, Hahn MW (2014) Reanalysis suggests that genomic islands of speciation are due to reduced diversity, not reduced gene flow. Molecular ecology, 23, 3133–57.

Cutter AD, Choi JY (2010) Natural selection shapes nucleotide polymorphism across the genome of the nematode Caenorhabditis briggsae Natural selection shapes nucleotide polymorphism across the genome of the nematode Caenorhabditis briggsae. Genome Research, 1103–1111.

Cutter AD, Payseur BA (2013) Genomic signatures of selection at linked sites: unifying the disparity among species. Nature reviews. Genetics, 14, 262–74.

Danecek P, Auton A, Abecasis G et al. (2011) The variant call format and VCFtools. Bioinformatics, 27, 2156–2158.

Denver DR, Dolan PC, Wilhelm LJ et al. (2009) A genome-wide view of Caenorhabditis elegans base-substitution mutation processes. Proceedings of the National Academy of Sciences of the United States of America, 106, 16310–4.

Dia I, Guelbeogo M, Ayala D (2013) Advances and Perspectives in the Study of the Malaria Mosquito Anopheles funestus. In: Anopheles mosquitoes - New insights into malaria vectors (ed Manguin S)

Dia I, Lochouarn L, Boccolini D, Costantini C, Fontenille D (2000) Spatial and temporal variations of the chromosomal inversion polymorphism of Anopheles funestus in Senegal. Parasite, 7, 179–184.

Dobzhansky T (1943) Genetics of natural populations. IX. Temporal changes in the composition of populations of Drosophila pseudoobscura. Genetics, 28, 162–186.

Dobzhansky T (1948) Genetics of natural populations .XVIII. Experiments on chromosomes of Drosophila pseudoobscura from different geographic regions. Genetics, 33, 588–602.

Dobzhansky T (1950) Genetics of natural populations. XIX. Origin of heterosis through natural selection in populations of Drosophila pseudoobscura. Genetics, 35, 288–302.

Dobzhansky T (1970) Genetics of the evolutionary process. New York.

Donnelly MJ, Licht MC, Lehmann T (2001) Evidence for recent population expansion in the evolutionary history of the malaria vectors Anopheles arabiensis and Anopheles gambiae. Molecular biology and evolution, 18, 1353–1364.

Earl DA, VonHoldt BM (2012) STRUCTURE HARVESTER: a website and program for visualizing STRUCTURE output and implementing the Evanno method. Conservation Genetics Resources, 4.

Edgley ML, Baillie DL, Riddle DL, Rose AM (2006) Genetic balancers (Wormbook, Ed,).

Evanno G, Goudet J, Regnaut S (2005) Detecting the number of clusters of individuals using the software structure: a simulation study. Molecular Ecology, 14, 2611–2620.

Excoffier L, Smouse PE, Quattro JM (1992) Analysis of molecular variance inferred from metric distances among DNA haplotypes: Application to human mitochondrial DNA restriction data. Genetics, 131, 479–491.

Feder JL, Egan SP, Nosil P (2012) The genomics of speciation-with-gene-flow. Trends in Genetics, 28, 342–350.

Felsenstein J (1974) The evolutionary advantage of recombination. II. Individual selection for recombination. Genetics, 78, 737–756.

Fouet C, Gray E, Besansky NJ, Costantini C (2012) Adaptation to aridity in the malaria mosquito Anopheles gambiae: chromosomal inversion polymorphism and body size influence resistance to desiccation. PloS one, 7, e34841.

Fouet C, Kamdem C, Gamez S, White BJ (2017) Extensive genetic diversity among populations of the malaria mosquito Anopheles moucheti revealed by population genomics. Infection, Genetics and Evolution, 48, 27–33.

Fumagalli M, Vieira FG, Linderoth T (2014) ngsTools: methods for population genetics analyses from Next-Generation Sequencing data. Bioinformatics.

Gattepaille LM, Jakobsson M, Blum MGB (2013) Inferring population size changes with sequence and SNP data: lessons from human bottlenecks. Heredity, 110, 409–19.

Gillespie JH (1997) Junk ain’t what junk does: Neutral alleles in a selected context. Gene, 205, 291–299.

Gillies MT, Coetzee M (1987) A supplement to the Anophelinae of Africa south of the Sahara. The South African Institute for Medical Research, Johannesburg.

Gillies MT, De Meillon B (1968) The Anophelinae of Africa South of the Sahara. Publications of the South African Institute for Medical Research, Johannesburg.

Giraldo-Calderon GI, Emrich SJ, MacCallum RM et al. (2015) VectorBase: an updated bioinformatics resource for invertebrate vectors and other organisms related with human diseases. Nucleic Acids Research, 43, D707–D713.

Gosset CC, Bierne N (2013) Differential introgression from a sister species explains high FST outlier loci within a mussel species. Journal of Evolutionary Biology, 26, 14–26.

Gray EM, Rocca K a C, Costantini C, Besansky NJ (2009) Inversion 2La is associated with enhanced desiccation resistance in Anopheles gambiae. Malaria journal, 8, 215.

Green CA, Hunt RH (1980) Interpretation of variation in ovarian polytene chromosomes of Anopheles funestus Giles, A. parensis Gillies, and A. aruni ? Genetica, 51, 187–195.

Haddrill PR, Halligan DL, Tomaras D, Charlesworth B (2007) Reduced efficacy of selection in regions of the Drosophila genome that lack crossing over. Genome biology, 8, R18.

Hellmann I, Ebersberger I, Ptak SE, Pääbo S, Przeworski M (2003) A Neutral Explanation for the Correlation of Diversity with Recombination Rates in Humans. The American Journal of Human Genetics, 72, 1527–1535.

Hellmann I, Prüfer K, Ji H et al. (2005) Why do human diversity levels vary at a megabase scale? Genome Research, 15, 1222–1231.

Hentges KE, Justice MJ (2004) Checks and balancers: balancer chromosomes to facilitate genome annotation. Trends in Genetics, 20.

Hill WG, Robertson A (1966) The effect of linkage on the limits of artificial selection. Genetical Research, 8, 269–294.

Hoffmann A a, Rieseberg LH (2008) Revisiting the Impact of Inversions in Evolution: From Population Genetic Markers to Drivers of Adaptive Shifts and Speciation? Annual review of ecology, evolution, and systematics, 39, 21–42.

Hoffmann AA, Sgro CM, Weeks AR (2004) Chromosomal inversion polymorphisms and adaptation. Trends in Ecology & Evolution, 19, 482–488.

Hoffmann AA, Weeks AR (2007) Climatic selection on genes and traits after a 100 year-old invasion: a critical look at the temperate-tropical clines in Drosophila melanogaster from eastern Australia. Genetica, 129, 133–147.

Huber CD, Nordborg M, Hermisson J, Hellmann I (2014) Keeping It Local: Evidence for Positive Selection in Swedish Arabidopsis thaliana. Molecular Biology and Evolution, 31, 3026–3039.

Hudson RR, Kaplan NL (1988) The coalescent process in models with selection and recombination. Genetics, 120, 819–829.

Jakobsson M, Rosenberg N (2007) CLUMPP: a cluster matching and permutation program for dealing with multimodality in analysis of population structure. Bioinformatics, 23, 1801–1806.

Jombart T (2008) adegenet: a R package for the multivariate analysis of genetic markers. Bioinformatics, 24, 1403–1405.

De Jong G, Bochdanovits Z (2003) Latitudinal clines in Drosophila melanogaster: body size, allozyme frequencies, inversion frequencies, and the insulin-signalling pathway. Journal of Genetics, 82, 207–223.

Joron M, Frezal L, Jones RT et al. (2011) Chromosomal rearrangements maintain a polymorphic supergene controlling butterfly mimicry. Nature, 477, 203–206.

Kamau L, Hunt RH, Coetzee M (2002) Analysis of the population structure of Anopheles funestus (Diptera: Culicidae) from western and coastal Kenya using paracentric chromosomal inversion frequencies. Journal of Medical Entomology, 39, 78–83.

Kamdem C, Fouet C, Gamez S, White BJ (2017) Pollutants and insecticides drive local adaptation in African malaria mosquitoes. Molecular Biology and Evolution, 34, 1261–1275.

Kaplan NL, Hudson RR, Langley CH (1989) The “hitchhiking effect” revisited. Genetics, 123, 887–899.

Kapun M, Van Schalkwyk H, McAllister B, Flatt T, Schlötterer C (2014) Inference of chromosomal inversion dynamics from Pool-Seq data in natural and laboratory populations of Drosophila melanogaster. Molecular Ecology, 23, 1813–1827.

Kapun M, Schmidt C, Durmaz E, Schmidt PS, Flatt T (2016) Parallel effects of the inversion In (3R) Payne on body size across the North American and Australian clines in Drosophila melanogaster. Journal of Evolutionary Biology, 29, 1059–1072.

Kawecki TJ, Dieter E (2004) Conceptual issues in local adaptation. Ecology Letters, 7, 1225–1241.

Keightley PD, Pinharanda A, Ness RW et al. (2015) Estimation of the Spontaneous Mutation Rate in Heliconius melpomene. Molecular Biology and Evolution, 32, 239–243.

Keinan A, Reich D (2010) Human population differentiation is strongly correlated with local recombination rate. PLoS Genetics, 6.

Kennington WJ, Hoffmann AA, Partridge L (2007) Mapping Regions Within Cosmopolitan Inversion In(3R)Payne Associated With Natural Variation in Body Size in Drosophila melanogaster. Genetics, 177, 549–556.

Kim Y, Maruki T (2011) Hitchhiking effect of a beneficial mutation spreading in a subdivided population. Genetics, 189, 213–226.

Kimura MK (1983) The Neutral Theory of Molecular Evolution. Cambridge, UK.

Kirkpatrick M (2010) How and why chromosome inversions evolve. PLoS Biology, 8.

Kirkpatrick M, Barton N (2006) Chromosome inversions, local adaptation and speciation. Genetics, 173, 419–434.

Kliman R, Hey J (1993) Reduced natural selection associated with low recombination in Drosophila melanogaster. Molecular biology and evolution, 10, 1239–1258.

Knibb WR (1982) Chromosome inversion polymorphisms in Drosophila melanogaster II. Geographic clines and climatic associations in Australasia, North America and Asia. Genetica, 58, 213–221.

Korneliussen T, Albrechtsen A, Nielsen R (2014) ANGSD: Analysis of Next Generation Sequencing Data. BMC Bioinformatics, 15, 356.

Korneliussen TS, Moltke I, Albrechtsen A, Nielsen R (2013) Calculation of Tajima’s D and other neutrality test statistics from low depth next-generation sequencing data. BMC bioinformatics, 14, 289.

Krimbas C, Powell JR (1992) Introduction. In: Drosophila inversion polymorphism (eds Krimbas C, Powell JR), p. London.

Kulathinal RJ, Bennett SM, Fitzpatrick CL, Noor MAF (2008) Fine-scale mapping of recombination rate in Drosophila refines its correlation to diversity and divergence. Proceedings of the National Academy of Sciences, 105, 10051–10056.

Kulathinal RJ, Stevison LS, Noor MAF (2009) The genomics of speciation in Drosophila: Diversity, divergence, and introgression estimated using low-coverage genome sequencing. PLoS Genetics, 5.

Lahn BT, Page DC (1999) Chromosome Four Evolutionary Strata on the Human X Chromosome. Science, 286, 964–967.

Lanzaro GC, Lee Y (2013) Speciation in Anopheles gambiae — The Distribution of Genetic Polymorphism and Patterns of Reproductive Isolation Among Natural Populations. In: Anopheles mosquitoes - New insights into malaria vectors (ed Manguin S)

Lee SF, Chen Y, Varan AK et al. (2011) Molecular Basis of Adaptive Shift in Body Size in Drosophila melanogaster: Functional and Sequence Analyses of the Dca Gene Research article. Molecular Biology and Evolution, 28, 2393–2402.

Lee Y, Meneses CR, Fofana A, Lanzaro GC (2009) Desiccation Resistance Among Subpopulations of Anopheles gambiae s.s. From Selinkenyi, Mali. Journal of medical entomology, 46, 316–320.

Lewontin RC, Krakauer J (1973) DISTRIBUTION OF GENE FREQUENCY AS A TEST OF THE THEORY O F THE SELECTIVE NEUTRALITY OF and LEWONTIN. Genetics, 74, 175–195.

Lischer HEL, Excoffier L (2012) PGDSpider: An automated data conversion tool for connecting population genetics and genomics programs. Bioinformatics, 28, 298–299.

Lowry DB, Willis JH (2010) A Widespread Chromosomal Inversion Polymorphism Contributes to a Major Life-History Transition, Local Adaptation, and Reproductive Isolation. PLoS Biology, 8.

Magni GE, Von Borstel RC (1962) Different Rates of Spontaneous Mutation during Mitosis and Meiosis in Yeast. Genetics, 47, 1097–1108.

Majewski J, Cohan FM (1999) Adapt globally, act locally: The effect of selective sweeps on bacterial sequence diversity. Genetics, 152, 1459–1474.

Maynard Smith J, Haigh J (1974) The hitch-hiking effect of a favourable gene. Genetical research, 23, 23–35.

Mcgaugh SE, Heil CSS, Manzano-winkler B et al. (2012) Recombination Modulates How Selection Affects Linked Sites in Drosophila. PLoS Biology, 10.

McGaugh SE, Noor M a F (2012) Genomic impacts of chromosomal inversions in parapatric Drosophila species. Philosophical transactions of the Royal Society of London. Series B, Biological sciences, 367, 422–9.

Meirmans P, Van Tienderen P (2004) GENOTYPE and GENODIVE: two programs for the analysis of genetic diversity of asexual organisms. Molecular Ecology Notes, 4, 792–794.

Mettler LE, Voelker RA, Mukai T (1977) Inversion clines in populations of Drosophila melanogaster. Genetics, 87, 169–176.

Michel a. P, Ingrasci MJ, Schemerhorn BJ et al. (2005) Rangewide population genetic structure of the African malaria vector Anopheles funestus. Molecular Ecology, 14, 4235–4248.

Muller HJ (1918) Genetic variability, twin hybrids, and constant hybrids, in a case of balanced lethal factors. Genetics, 3, 422–499.

Nachman MW (2002) Variation in recombination rate across the genome: Evidence and implications. Current Opinion in Genetics and Development, 12, 657–663.

Nachman MW, Payseur BA (2012) Recombination rate variation and speciation: theoretical predictions and empirical results from rabbits and mice. Philosophical Transactions of the Royal Society B: Biological Sciences, 367, 409–421.

Neafsey DE, Waterhouse RM, Abai MR et al. (2015) Highly evolvable malaria vectors: The genomes of 16 Anopheles mosquitoes. Science, 347, 1258522–1258522.

Nielsen R (2005) Molecular Signatures of Natural Selection. Annual review of genetics, 39, 197–218.

Nishikawa H, Iijima T, Kajitani R et al. (2015) A genetic mechanism for female-limited Batesian mimicry in Papilio butterfly. Nature genetics, 47, 405–9.

Noor MAF (2008) Mutagenesis from meiotic recombination is not a primary driver of sequence divergence between Saccharomyces species. Molecular Biology and Evolution, 25, 2439–2444.

Noor M a F, Bennett SM (2009) Islands of speciation or mirages in the desert? Examining the role of restricted recombination in maintaining species. Heredity, 103, 439–444.

Noor MAF, Grams KL, Bertucci LA, Reiland J (2001) Chromosomal inversions and the reproductive isolation of species. Proceedings of the National Academy of Sciences, 98, 12084–12088.

Nordborg M, Charlesworth B, Charlesworth B (1996) Increased levels of polymorphism surrounding selectively maintained sites in highly selfing species. Proceedings of the Royal Society B: Biological Sciences, 263, 1033–1039.

Nordborg M, Hu TT, Ishino Y et al. (2005) The pattern of polymorphism in Arabidopsis thaliana. PLoS Biology, 3, 1289–1299.

Nordborg M, Tavar S (2002) Linkage disequilibrium: what history has to tell us. Trends in Genetics, 18, 83–90.

Nosil P, Feder JL (2012) Widespread yet heterogeneous genomic divergence. Molecular Ecology, 21, 2829–2832.

Nosil P, Funk DJ, Ortiz-Barrientos D (2009) Divergent selection and heterogeneous genomic divergence. Molecular Ecology, 18, 375–402.

O’Loughlin SM, Magesa S, Mbogo C et al. (2014) Genomic Analyses of Three Malaria Vectors Reveals Extensive Shared Polymorphism but Contrasting Population Histories. Molecular biology and evolution, 1–14.

Ortiz-barrientos D, Engelstädter J, Rieseberg LH (2016) Recombination Rate Evolution and the Origin of Species. Trends in Ecology & Evolution, 31, 226–236.

Ortiz-Barrientos D, Reiland J, Hey J, Noor MAF (2002) Recombination and the divergence of hybridizing species. Genetica, 116, 167–178.

Paradis E, Claude J, Strimmer K (2004) Analyses of Phylogenetics and Evolution in R language. Bioinformatics, 20, 289–290.

Pegueroles C, Ordóñez V, Mestres F, Pascual M (2010) Recombination and selection in the maintenance of the adaptive value of inversions. Journal of Evolutionary Biology, 23, 2709–2717.

Peterson BK, Weber JN, Kay EH, Fisher HS, Hoekstra HE (2012) Double Digest RADseq: An Inexpensive Method for De Novo SNP Discovery and Genotyping in Model and Non-Model Species. PLoS ONE, 7, e37135.

Pool JE, Corbett-Detig RB, Sugino RP et al. (2012) Population Genomics of Sub-Saharan Drosophila melanogaster: African Diversity and Non-African Admixture. PLoS Genetics, 8.

Powell JR (1997) Progress and prospects in evolutionary biology: the Drosophila model. New York.

Pritchard JK, Stephens M, Donnelly P (2000) Inference of population structure using multilocus genotype data. Genetics, 155, 945–959.

Purcell S, Neale B, Todd-Brown K et al. (2007) PLINK: a toolset for whole-genome association and population-based linkage analysis. American Journal of Human Genetics, 81.

R Development Core Team (2014) R: A language and environment for statistical computing. R Foundation for Statistical Computing, Vienna, Austria.

Rako L, Anderson AR, Sgro CM, Stocker AJ, Hoffmann AA (2006) The association between inversion In (3R) Payne and clinally varying traits in Drosophila melanogaster. Genetica, 128, 373–384.

Riehle MM, Guelbeogo WM, Gneme A et al. (2011) A cryptic subgroup of Anopheles gambiae is highly susceptible to human malaria parasites. Science (New York, N.Y.), 331, 596–8.

Rieseberg LH (2001) Chromosomal rearrangements and speciation. Trends in Ecology and Evolution, 16, 351–358.

Roberts P (1976) The genetics of chromosomal aberration. In: The genetics and biology of Drosophila (eds Ashburner M, Novitski E), pp. 67–184. London.

Roesti M, Hendry AP, Salzburger W, Berner D (2012) Genome divergence during evolutionary diversification as revealed in replicate lake-stream stickleback population pairs. Molecular ecology, 21, 2852–62.

Roesti M, Moser D, Berner D (2013) Recombination in the threespine stickleback genome - Patterns and consequences. Molecular Ecology, 22, 3014–3027.

Rosenberg N (2004) DISTRUCT: a program for the graphical display of population structure. Molecular Ecology Resources, 4, 137–138.

Savolainen O, Lascoux M, Merilä J (2013) Ecological genomics of local adaptation., 14.

Schaeffer SW (2008) Selection in heterogeneous environments maintains the gene arrangement polymorphism of drosophila pseudoobscura. Evolution, 62, 3082–3099.

Schaeffer SW, Goetting-minesky MP, Kovacevic M et al. (2003) Evolutionary genomics of inversions in Drosophila pseudoobscura: Evidence for epistasis. Proceedings of the National Academy of Sciences, 100, 8319–8324.

Schlötterer C (2002) Towards a molecular characterization of adaptation in local populations. Current Opinion in Genetics and Development, 12, 683–687.

Schneider KA, Kim Y (2013) Genetic Hitchhiking under Heterogeneous Spatial Selection Pressures. PLoS ONE, 8.

Seehausen O, Butlin RK, Keller I et al. (2014) Genomics and the origin of species. Nature reviews. Genetics, 15, 176–92.

Service MW (1993) Mosquito ecology: field sampling methods (UK Elsevier Applied Science, London, Ed,).

Sharakhova M V, Xia A, Leman SC, Sharakhov I V (2011) Arm-specific dynamics of chromosome evolution in malaria mosquitoes. BMC evolutionary biology, 11, 91.

Sharakhov I, Braginets O, Grushko O et al. (2004) A Microsatellite Map of the African Human Malaria Vector Anopheles funestus. Journal of Heredity, 95, 29–34.

Sinka ME, Bangs MJ, Manguin S et al. (2010) The dominant Anopheles vectors of human malaria in Africa, Europe and the Middle East: occurrence data, distribution maps and bionomic précis. Parasites & vectors, 3, 117.

Slatkin M, Wiehe T (1998) Genetic hitch-hiking in a subdivided population. Genetical Research, 71, 155–160.

Spencer CCA, Deloukas P, Hunt S et al. (2006) The influence of recombination on human genetic diversity. PLoS Genetics, 2, 1375–1385.

Stephan W, Langley CH (1989) Molecular genetic variation in the centromeric region of the X chromosome in three Drosophila ananassae populations. I. Contrasts between the vermilion and forked loci. Genetics, 121, 89–99.

Storz JF (2005) Using genome scans of DNA polymorphism to infer adaptive population divergence. Molecular Ecology, 14, 671–688.

Stump AD, Fitzpatrick MC, Lobo NF et al. (2005) Centromere-proximal differentiation and speciation in Anopheles gambiae. Proceedings of the National Academy of Sciences of the United States of America, 102, 15930–5.

Sturtevant A (1921) A case of rearrangement of genes in Drosophila. Proceedings of the National Academy of Sciences, 7, 235–237.

Tajima F (1989) Statistical method for testing the neutral mutation hypothesis by DNA polymorphism. Genetics, 123, 585–595.

Tenaillon MI, U’Ren J, Tenaillon O, Gaut BS (2004) Selection versus demography: A multilocus investigation of the domestication process in maize. Molecular Biology and Evolution, 21, 1214–1225.

Thomas CG, Wang W, Jovelin R et al. (2015) Full-genome evolutionary histories of selfing, splitting, and selection in Caenorhabditis., 667–678.

Thornton K (2005) Recombination and the properties of Tajima’s D in the context of approximate-likelihood calculation. Genetics, 171, 2143–8.

Tiffin P, Ross-Ibarra J (2014) Advances and limits of using population genetics to understand local adaptation. Trends in Ecology & Evolution, 29, 673–680.

Watterson GA (1975) On the number of segregating sites in genetical models without recombination. Theoretical Population Biology, 7, 256–276.

Weir BS, Cockerham CC (1984) Estimating F-statistics for the analysis of population structure. Evolution, 38, 1358–1370.

Williams GC (1966) Adaptation and Natural Selection: A Critique of Some Current Evolutionary Thought. Princeton, NJ.

Wright S, Dobzhansky T (1946) Genetics of natural populations. XII. Experimental reproduction of some of the changes caused by natural selection in certain populations of Drosophila pseudoobscura. Genetics, 31, 125–156.

Wright SI, Foxe JP, DeRose-Wilson L et al. (2006) Testing for effects of recombination rate on nucleotide diversity in natural populations of Arabidopsis lyrata. Genetics, 174, 1421–1430.

Wu TD, Nacu S (2010) Fast and SNP-tolerant detection of complex variants and splicing in short reads. Bioinformatics, 26, 873–881.

Yeaman S (2013) Genomic rearrangements and the evolution of clusters of locally adaptive loci. Proceedings of the National Academy of Sciences, 110, E1743-51.

